# A multifaceted suite of metrics for comparative myoelectric prosthesis controller research

**DOI:** 10.1101/2023.08.28.554998

**Authors:** Heather E. Williams, Ahmed W. Shehata, Kodi Y. Cheng, Jacqueline S. Hebert, Patrick M. Pilarski

## Abstract

Upper limb robotic (myoelectric) prostheses are technologically advanced, but challenging to use. In response, substantial research is being done to develop user-specific prosthesis controllers that can predict a person’s intended movements. Most studies that test and compare new controllers rely on simple assessment measures such as task scores (e.g., number of objects moved across a barrier) or duration-based measures (e.g., overall task completion time). These assessment measures, however, fail to capture valuable details about: the quality of device arm movements; whether these movements match users’ intentions; the timing of specific wrist and hand control functions; and users’ opinions regarding overall device reliability and controller training requirements. In this work, we present a comprehensive and novel suite of myoelectric prosthesis control evaluation metrics that better facilitates analysis of device movement details—spanning measures of task performance, control characteristics, and user experience. As a case example of their use and research viability, we applied these metrics in real-time control experimentation. Here, eight participants without upper limb impairment compared device control offered by a deep learning-based controller (recurrent convolutional neural network-based classification with transfer learning, or RCNN-TL) to that of a commonly used controller (linear discriminant analysis, or LDA). The participants wore a simulated prosthesis and performed complex functional tasks across multiple limb positions. Analysis resulting from our suite of metrics identified 16 instances of a user-facing problem known as the “limb position effect”. We determined that RCNN-TL performed the same as or significantly better than LDA in four such problem instances. We also confirmed that transfer learning can minimize user training burden. Overall, this study contributes a multifaceted new suite of control evaluation metrics, along with a guide to their application, for use in research and testing of myoelectric controllers today, and potentially for use in broader rehabilitation technologies of the future.

## Introduction

Below elbow (transradial) is the most prevalent of major upper limb amputations [1]. A myoelectric prosthesis offers a means of restoring complex limb function to those with transradial amputation, ideally across a wide range of arm positions [2]. Conventional myoelectric device control is based on electromyography (EMG) [3]. Here, signals are typically detected by surface electrodes that are housed within a donned prosthesis socket and then transmitted to the device’s onboard controller. The controller decodes user-specific muscle contractions and sends corresponding instructions to appropriate prosthesis wrist and hand motors.

Myoelectric prostheses that employ pattern recognition offer predictive device control that is capable of learning a user’s intended movements [4, 5]. Despite the potential of such machine learning-based control solutions, device performance challenges persist for users, particularly when various limb positions are necessary [6]. In these instances, EMG signals change due to gravity, supplemental muscle activities, and electrode shifts resulting from changes in muscle topology [7]. Control in such instances can be unpredictable and therefore frustrating for users [6]. This control challenge is well documented and referred to as the “limb position effect” [7]. Several pattern recognition-based control methods have been investigated to minimize the limb position effect [8–22]. These methods require a user to execute a training routine across multiple limb positions, prior to daily device use. A training routine involves execution of a specific sequence of forearm muscle contractions. EMG signals resulting from the muscle contractions are captured for use by the device controller’s model. The model learns to recognize various patterns of consistent and repeated muscle signal features [4], including patterns involved in wrist rotation and hand open/close. Learned features are subsequently classified during device use, and the resulting classifications inform device motor instructions.

Inertial measurement unit (IMU) data can provide a classification control model with additional and informative limb position-related data [7, 23]. Deep learning control methods, such as recurrent convolutional neural networks (RCNNs), can combine high volumes of EMG and IMU data from multiple limb positions. However, to capture all required muscle and limb position data (in low to high arm positions), lengthy and burdensome training routines must be performed by users [2, 24, 25]. Control model retraining is also required in instances when device control degrades, such as due to muscle fatigue or electrode shifts. The overall training burden poses drawbacks to position-aware myoelectric control methods.

Our earlier study uncovered that the addition of Transfer Learning (TL) can alleviate the training burden necessitated by data intensive RCNN-based solutions [26]. In this previous work, an RCNN classification control model (classifier) was trained using a large dataset of EMG and IMU signals obtained from numerous individuals with intact upper limbs, to become the starting point of new users’ control (with a simulated prosthesis donned [27, 28]). Each new user required a reduced amount of personal data for training thereafter. Not only did this research confirm that RCNN-based classification control with TL reduces training burden, but it also offered a prosthesis control solution with a tendency towards better functional task performance across multiple limb positions, when compared to a linear discriminant analysis (LDA) classification controller. Interestingly, the research identified possible instances of the limb position effect during high grasping movements, however it was noted that more detailed measures of control were needed to confirm this [26]. As a corollary to the TL-based findings resulting from this work, metrics deficiencies were uncovered—control characteristics outcomes evidenced during task performance could not be fully understood, and user-reported control experiences were not considered [26].

There remain gaps in the completeness of metrics used for prosthesis control research. Most studies that test myoelectric control via functional tasks, tend only to examine general performance metrics, including task success rates or duration-based measures [29–37]. These metrics, however, cannot yield a complete understanding of the quality of participants’ hand, wrist, and arm movements [38]. Furthermore, they cannot adequately characterize the nature of device control, such as the identification of unnecessary grip aperture adjustments [26]. For these reasons, our earlier work recommended that RCNN-based classification control with TL (RCNN-TL) not be judged by task performance alone, but rather, that control characteristics also be considered. Then collectively, the task performance and control characteristics can be weighed against subjective user experience, to provide a full complement of data-driven prosthesis control outcomes.

This current study contributes a comprehensive suite of metrics that aims to address the issue of inconclusive myoelectric controller assessment outcomes. The suite includes three broad categories of metrics: **task performance**, **control characteristics**, and **user experience**. We deployed these metrics, as part of this current work, to reinvestigate our earlier controller research findings [26]—examining whether RCNN-TL can indeed reduce training burden, offer improved device control over a comparative LDA baseline classification controller (LDA-Baseline), and if instances of the limb position effect can be pinpointed. In using the novel suite of metrics, this work contributes a data-driven understanding of *when* and *why* TL-based neural network control solutions show great promise towards solving the limb position effect challenge. It is expected that the suite of evaluation metrics introduced by this research will guide future rehabilitation device control experimentation.

What follows is a presentation of our metrics, an overview of our reinvestigation research, and a detailed presentation of its methods, results, discussion, and conclusion.

## Introducing the Suite of Myoelectric Control Evaluation Metrics

Performance-based assessments for the evaluation of real-time upper limb prosthesis control often require participants to execute on-screen virtual arm movements or on-screen cursor movements (such as the Target Achievement Control test [39] and Fitts’ Law tests [40], respectively) [8, 14, 41–51]. EMG sensors placed on participants’ limbs record muscle activations in such assessments. However, limb kinematics and factors that change EMG signals (including the limb position effect) are not taken into consideration [44]. Consequently, research that employs on-screen assessments often recommend that future work be undertaken using alternative real-time methods [8, 43].

Other studies have taken a next step towards a deeper understanding of control, through the introduction of functional task assessments [29–37, 51–53]. Here, either non-disabled participants wearing a simulated prosthesis [27] or actual myoelectric prosthesis users, are required to perform upper limb tasks that mimic activities of daily living. The movement of participants’ upper limbs during task execution can be recorded using motion capture technology. From the resulting data, hand movement metrics, including hand velocity, hand distance travelled, and hand trajectory variability can be calculated [38, 54–56]. We used these hand movement metrics, plus common task success rate and duration-based measures, and present them as **Task Performance Metrics**. All such metrics can be found in Table 1, with calculations derived from the work of Valevicius et al. [38]. S1 Table offers select examples of strong and weak outcomes resulting from these metrics.

**Table 1.** Description of Task Performance metrics used in analysis. For each metric, the following details are outlined: the metric’s name; a description of the metric; the data analysis level at which the metric is calculated (Controller, Task, Trial, Movement, Movement Segment, or Phase); the data required for the metric calculation; the metric calculation procedure; and indicators that constitute a control improvement.

| Metric | Metric Description | Metric Calculation |  |  | Control Improvement Indicator<br>(∴ means because) |
| --- | --- | --- | --- | --- | --- |
|  |  | Data Analysis Level | Required Data | Calculation Procedure |  |
| Success Rate | Percent of trials that are error-free (%) | Task | Number of error-free trials | The number of error-free trials divided by the number of possible trial attempts [35] | A higher success rate ∴ participants make fewer errors |
| Trial Duration | Elapsed time for each trial (sec) | Trial | Timestamps | The timestamp at the end of the trial's last phase minus the timestamp at the start of trial's first phase [35] | A shorter trial duration ∴ participants complete trials quicker, also indicative of higher skill level [25], [55] |
| Phase Duration | Elapsed time for each phase (sec) | Phase: Reach, Grasp, Transport, Release | Timestamps | The timestamp at the end of the phase minus the timestamp at the beginning of the phase [35] | A shorter phase duration ∴ participants complete phases quicker, also indicative of higher skill level [25], [55] |
| Relative Phase Duration | Elapsed time for each phase, relative to the elapsed time for a Reach-Grasp-Transport-Release movement (%) | Phase: Reach, Grasp, Transport, Release | Timestamps | The phase duration divided by the movement duration (i.e., the difference between the timestamps at the beginning and end of a Reach-Grasp-Transport-Release movement) [35] | For Grasp and Release phases: A shorter relative phase duration ∴ participants manipulate objects with more ease [25]<br>For Reach and Transport phases: Shorter Grasp and Release relative phase durations lead to longer Reach and Transport relative phase durations [25] |
| Peak Hand Velocity | Maximum velocity of the hand while moving (mm/s) | Movement Segment: Reach-Grasp, Transport-Release | 3D Position of the Hand (mm, mm, mm) | The maximum hand velocity achieved within a movement segment [35] | A higher peak hand velocity ∴ participants move their hand quicker |
| Hand Distance Travelled | Total distance travelled by the hand while moving (mm) | Movement Segment: Reach-Grasp, Transport-Release | 3D Position of the Hand (mm, mm, mm) | Calculation steps:<br>1. The distance between the 3D positions of the hand at successive sampled points in time are calculated<br>2. The sum of these distances is calculated [35] | A shorter hand distance travelled ∴ the hand movement paths of participants are more efficient [25] |
| Hand Trajectory Variability | How much the hand movement path varies between trials (mm) | Movement Segment: Reach-Grasp, Transport-Release<br>(Note that hand trajectory variability is calculated for each participant, rather than for each trial) | 3D Position of the Hand (mm, mm, mm) | Calculation steps, used for each participant:<br>1. The 3D position (x, y, z) of the hand is time-normalized to 101 sampled points in time (0%, 1%, ..., 100% of the movement segment) for each of the participant's trials<br>2. At each sampled point, the standard deviation of the x, y, and z positions across the participant's trials are independently calculated<br>3. The mean of the resulting standard deviations is calculated at each sampled point<br>4. The maximum of these means is determined [35] | A smaller hand trajectory variability ∴ participants do not explore a variety of hand movement paths<br>(possibly indicating confidence in their chosen path with movements in keeping with those of unimpaired individuals) [25], [35] |

Task performance metrics alone, however, do not capture the nuances of device control characteristics, such as instances where a user introduces unnecessary hand/wrist movements when grasping or releasing an object [26]. Some studies have introduced metrics that quantify specific myoelectric prosthesis control characteristics, including misclassification rates [57], grasp force [57, 58], grip aperture plateau time [59], wrist rotation range of motion [57], workload (assessed via pupil size) [60], and measures of muscle activations [61]. We selected and derived metrics from the literature, plus developed additional novel metrics, and collectively present them as **Control Characteristic Metrics**. All such metrics can be found in Table 2, with select examples in S1 Table.

**Table 2.** Description of Control Characteristics metrics used in analysis. For each metric, the following details are outlined: the metric’s name; a description of the metric; the data analysis level at which the metric is calculated (Controller, Task, Trial, Movement, Movement Segment, or Phase); the data required for the metric calculation; the metric calculation procedure; and indicators that constitute a control improvement.

| Metric | Metric Description | Metric Calculation |  |  | Control Improvement Indicator<br>(∴ means because) |
| --- | --- | --- | --- | --- | --- |
|  |  | Data Analysis Level | Required Data | Calculation Procedure |  |
| Total Grip Aperture Movement | Total amount of grip aperture variation (mm) | *Phase:*<br>Reach, Grasp, Transport, Release | Grip Aperture (mm) | 1. The absolute differences between the grip apertures at successive sampled points in time are calculated<br>2. The sum of these differences is calculated | A smaller total grip aperture movement ∴ fewer unnecessary hand open/close movements occur |
| Total Wrist Rotation Movement | Total amount of wrist rotation angle variation (deg) | *Phase:*<br>Reach, Grasp, Transport, Release | Wrist Rotation Angle (degree) | 1. The absolute differences between the wrist rotation angles at successive sampled points in time are calculated<br>2. The sum of these differences is calculated | A smaller total wrist rotation movement ∴ fewer unnecessary wrist rotation movements occur |
| Number of Grip Aperture Adjustments | Number of times that grip aperture variation commences or changes direction | *Phase:*<br>Reach, Grasp, Transport, Release<br>(Note that Transport adjustments may not cause object drop) | Grip Aperture (mm) | 1. The grip aperture velocity is smoothed using a moving average filter<br>2. The sign of the smoothed velocity is used to detect the following: a change from 0 to positive; a change from 0 to negative; a change from negative to positive; a change from positive to negative<br>3. The number of times that the above sign changes occurred is tallied [61] | A smaller number of grip aperture adjustments ∴ fewer misclassifications are made [61] |
| Number of Wrist Rotation Adjustments | Number of times that wrist rotation angle variation commences or changes direction | *Phase:*<br>Reach, Grasp, Transport, Release | Wrist Rotation Angle (degree) | The number of wrist rotation adjustments were calculated as per grip aperture velocity detailed above (but with wrist rotation data rather than grip aperture data). | A smaller number of wrist rotation adjustments ∴ fewer misclassifications are made [61] |
| Grip Aperture Plateau | Amount of time during which the grip aperture remains open before closing to grasp a task object (sec) | *Movement Segment:*<br>Reach-Grasp | Grip Aperture (mm) | 1. The grip aperture and grip aperture velocity are both smoothed using a moving average filter<br>2. Plateaus are identified when both of the following conditions are met:<br>- the smoothed grip aperture is greater than 90% of the maximum grip aperture in the movement segment<br>- the smoothed grip aperture velocity is less than 10% of the maximum grip aperture velocity in the movement segment<br>3. The sum of the time duration(s) in which the above conditions are met is calculated<br>(modified from Bouwsema et al.'s calculation [55]) | A shorter plateau ∴ more natural movements are made (more closely mimicking those of unimpaired individuals) and higher skill level is evident [25], [55] |
| Simultaneous Wrist-Shoulder Movements | Percent of the phase during which the wrist rotation is controlled while the shoulder is moving (%) | *Phase:* Any phase that requires both wrist rotation and shoulder movements. | Wrist Rotation Angle (degree) & Shoulder Flexion/Extension Angle (degree) | 1. Wrist rotation and shoulder flexion/extension angular velocities are smoothed using a moving average filter<br>2. The absolute values of the smoothed angular velocities are calculated<br>3. Simultaneous movements are identified when both of the following conditions are met:<br>- The smoothed wrist rotation angular velocity is above a threshold of 10% of the maximum wrist rotation angular velocity in the movement<br>- The smoothed shoulder flexion/extension angular velocity is above a threshold of 10% of the maximum shoulder flexion/extension angular velocity in the movement<br>4. The sum of the time duration(s) in which the above conditions are met is calculated<br>5. The resulting sum is divided by the phase duration (i.e., the timestamp at the end of the phase minus the timestamp at the beginning of the phase) | A larger percent of simultaneous wrist-shoulder movements ∴ participants control wrist rotation while moving their arm, rather than pausing arm movements to focus on device control—in keeping with unimpaired movements |
| Total Muscle Activity | Total amount of muscle activity expended | *Phase:*<br>Reach, Grasp, Transport, Release | EMG signals (range of -1 to 1) | 1. The EMG signals are filtered and rectified<br>2. Each EMG channel is normalized using training data: a) signals from the rest wrist position with the elbow bent at 90° are obtained from that session's training data; b) step 2a rest signals are rectified; c) the mean of the step 2b rectified signals is calculated for each channel; d) each channel's mean is subtracted from that channel's step 1 signals<br>3. Step 2 signals are summed across channels<br>4. The envelope of the step 3 signals is obtained<br>5. The step 4 envelope is summed across time (similar to the calculation of Ingraham et al.'s composite sum EMG metric [57]) | A smaller value ∴ participants exert less muscle activity |

Finally, whether any proposed control solution yields noticeable improvement depends on users’ assessments. The National Aeronautics and Space Administration Task Load Index (NASA-TLX) is a survey tool, which measures subjective mental workload [62]. A recent literature review confirmed that the NASA-TLX has been widely employed in prosthesis use assessments [63]. Usability surveys offer yet another assessment approach and capture the users’ opinions of alternative device control solutions [64]. We selected relevant survey questions from the literature and present them as **User Experience Metrics**. All such metrics can be found in Table 3, with select examples in S1 Table.

**Table 3.** Description of User Experience metrics used in analysis. For each metric, the following details are outlined: the metric’s name; a description of the metric; the data analysis level at which the metric is calculated (Controller, Task, Trial, Movement, Movement Segment, or Phase); the data required for the metric calculation; the survey question; and indicators that constitute a control improvement.

| Metric | Metric Description |  | Metric Calculation |  |  | Control Improvement Indicator<br>(∴ means because) |
| --- | --- | --- | --- | --- | --- | --- |
|  |  |  | Data Analysis Level | Required Data | Survey Question |  |
| NASA-TLX | Mental Demand | Workload demand resulting from each controller | Controller | Survey Scores (rating 0–100) | How much mental and perceptual activity was required (e.g., thinking, remembering, looking, etc.)? Was the control easy or demanding, simple or complex? [58] | A smaller score ∴ participants experience less workload demand |
|  | Physical Demand |  |  |  | How much physical activity was required (e.g., very strong muscle contractions, upper arm movements, trunk movements, etc.)? Was the control easy or demanding, restful or laborious? [58] |  |
|  | Temporal Demand |  |  |  | How much time pressure did you feel to complete the tasks with this control strategy? Was the pace slow and leisurely or rapid and frantic? [58] |  |
|  | Performance |  |  |  | How hard did you have to work (mentally & physically) to accomplish your level of performance? [58] |  |
|  | Effort |  |  |  | How successful do you think you were in accomplishing the goals of the task with this control strategy? How satisfied were you with your performance in accomplishing these goals? [58] |  |
|  | Frustration |  |  |  | How insecure, discouraged, irritated, stressed, and/or annoyed versus secure, gratified, content, relaxed, and/or complacent did you feel during the task? [58] |  |
| Usability Survey | Intuitiveness | Usability of each controller | Controller | Survey Scores (rating 0–5) | Intuitiveness is defined as how easy it was to learn how to use the controller.<br>How easy is the controller to learn? [60] | A larger score ∴ participants report greater ease of use (device controllability) |
|  | Task Effectiveness |  |  |  | Effectiveness is defined as how well the controller was able to perform the task.<br>How well did the controller perform the task? [60] (asked for each task performed) |  |
|  | Reliability |  |  |  | Reliability is defined as how often the controller did something that was intended or expected.<br>How often did the arm move in a way that you wanted or expected? [60] |  |

Tables 1, 2, and 3, collectively describe the **Suite of Myoelectric Control Evaluation Metrics** introduced in this work. The Control Improvement Indicator column in each of these tables uses unimpaired limb movement as a yardstick for control assessment. To conduct such assessments, data collection protocols should include the following:

- Control models under investigation should be trained by participants using a research-specific series of hand/wrist movements that elicit forearm muscle signals for capture
- Participants must either wear a myoelectric prosthesis or a simulated prosthesis
- Participants must perform functional task(s) that can be split into the distinct phases of Reaches, Grasps, Transports, and Releases
- Motion capture data, muscle signals, and device motor data should be collected during functional task execution
- User experience survey responses should be collected at the end of each testing session
- Data streams of interest (for each functional task trial) could include: the number of error-free trials executed; trial time stamps; the 3D position of the device hand, its grip aperture and wrist rotation angles; plus the participants’ shoulder flexion/extension angles, EMG signal data, and post-testing session survey scores.

With adherence to the above-mentioned data collection requirements, the suite of control evaluation metrics presented in this work facilitates in-depth analysis that will uncover numerous upper limb prosthesis control insights. These insights are expected to be particularly beneficial in investigations that compare myoelectric device controllers.

## Overview: Reinvestigation using our Suite of Metrics

As an example of how the suite of evaluation metrics introduced in this work can be used to advance prosthesis control research, we deployed them in a deliberately challenging experiment—reinvestigating our earlier comparative classifier research findings [26]. Here, device control offered by two classifiers was compared: a proposed deep learning-based controller (RCNN-TL) intent on mitigating the limb position effect, versus a commonly used and commercially available controller (LDA-Baseline). Fig 1 presents an overview of how each control model was trained and tested, using two distinct groups of participants without upper limb impairment who wore an EMG and IMU data capture armband.

1. A large General Participant Group’s data created a control starting point for new users. **RCNN-TL’s Model Pre-Training**—Each General Participant Group member performed a training routine (isotonic forearm muscle contractions were executed in four limb positions), during which their forearm EMG and IMU signals were collected. Their collective resulting signal data, along with the corresponding classes of muscle contractions, informed RCNN-TL’s pre-trained control model.
2. A new, smaller Simulated Prosthesis (SP) Participant Group wore a simulated prosthesis. **RCNN-TL’s Model Retraining & Testing**—Each SP Participant Group member performed a brief training routine (isometric contractions were held in three limb positions). The resulting participant-specific EMG and IMU data, plus classes of muscle contractions, were subsequently used to calibrate RCNN-TL’s model. Participants tested RCNN-TL by performing functional tasks across multiple limb positions—the Pasta Box Task [38] and the Refined Clothespin Relocation Test (RCRT) [65]. **LDA-Baseline’s Model Training & Testing**—The forearm muscle signals of each SP Participant Group member were also captured using a standard pattern recognition training routine that was not designed to mitigate the limb position effect (isometric contractions were held in one limb position) [23]. The resulting EMG data, plus classes of muscle contractions, were used to train LDA-Baseline’s model. Each SP Participant Group member tested LDA-Baseline by performing the Pasta Box Task [38] and RCRT [65].

**Fig 1.**
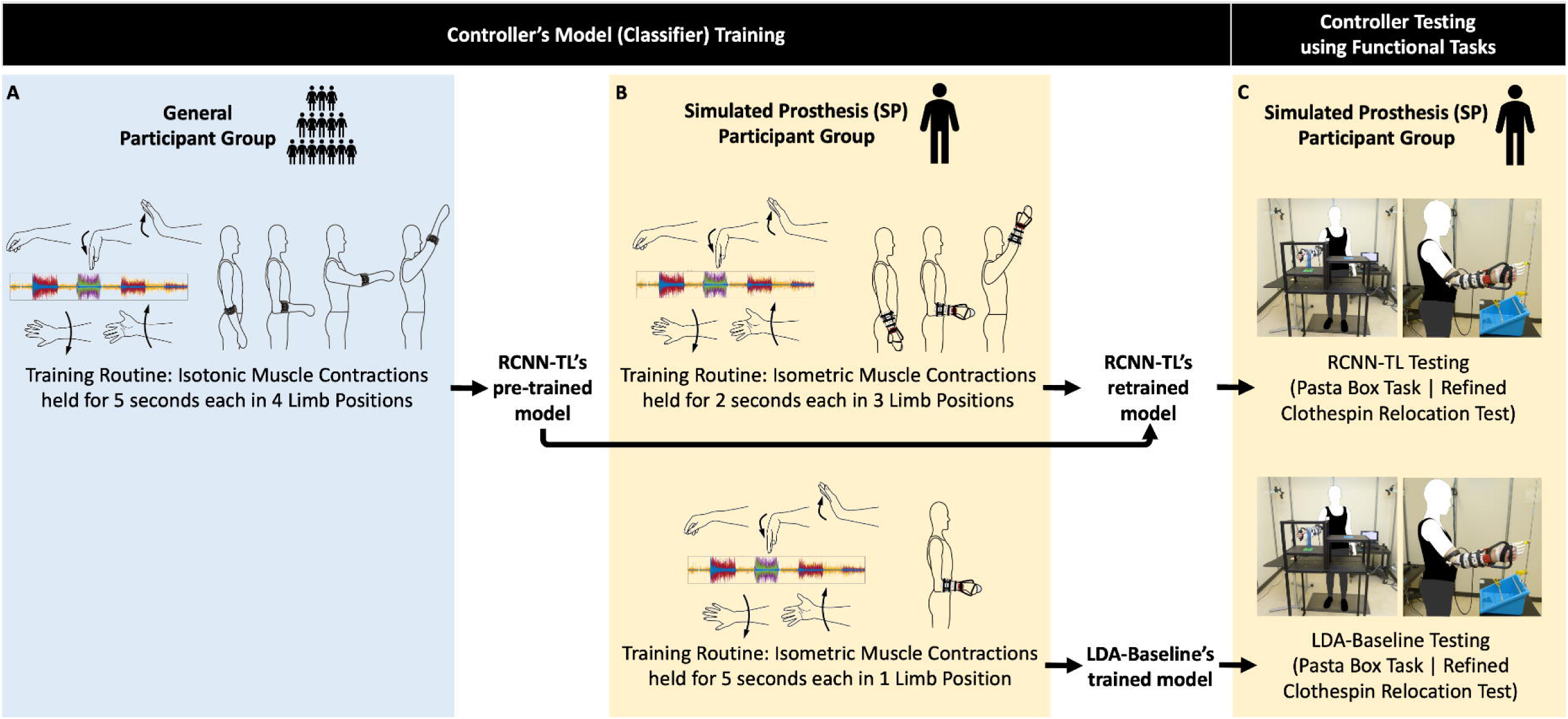
RCNN-TL and LDA-Baseline’s model training and testing. The blue panel (A) illustrates the step that the General Participant Group performed (training routine that yielded RCNN-TL’s pre-trained model) while wearing an EMG and IMU armband. The yellow panels illustrate the steps that the Simulated Prosthesis (SP) Participant Group performed: (B) respective training routines that yielded RCNN-TL’s retrained model and LDA-Baseline’s trained model, and (C) subsequent controller testing using functional tasks, all while wearing an EMG and IMU armband plus a simulated prosthesis.

## Methods

What follows are details about our reinvestigation research methods, including: participant descriptors; muscle signal data collection and processing techniques; a description of the simulated prosthesis donned by participants; specifications of the control models under investigation and their training requirements; setup of the testing environment; the functional tasks used to assess control; the participant survey administration process; control data processing techniques to yield the suite of metrics; the statistical analysis of such metrics; and the identification of instances of the limb position effect resulting from this analysis.

### Participants

Participants recruitment took place from March 2, 2022, to March 31, 2022. All participants provided written informed consent, as approved by the University of Alberta Health Research Ethics Board (Pro00086557).

**General Participant Group** *(without simulated prosthesis)* —Nineteen participants without upper limb impairment were recruited. All had normal or corrected vision, 10 were male, nine were female, 17 were right-handed. They had a median age of 25 years (range: 19–58 years) and median height of 170 cm (range: 159–193 cm). Each of the 19 participants completed one data collection session.

**SP Participant Group** *(with donned simulated prosthesis)* —A total of nine new participants without upper limb impairment were recruited. One participant was removed due to their inability to reliably control the donned simulated prosthesis even after control practice. Of the remaining eight participants, all had normal or corrected vision, five were male, three were female, seven were right-handed. They had a median age of 22 years (range: 20–56 years) and median height of 181 cm (range: 169–185 cm). No participants had experience with EMG pattern recognition control using a simulated prosthesis. The eight participants completed two data collection sessions on different days, with a median of 24 days between sessions (range: 18 – 45 days). Half of the participants retrained/tested RCNN-TL in their first session (as shown in Fig 1B–C), and the other half trained/tested LDA-Baseline in their first session (also shown in Fig 1B–C). Each trained/tested the other controller in their second session.

### Signal Data Collection & Processing Procedure

Participants in both groups wore a Myo gesture control armband (Thalmic Labs, Kitchener, Canada)over their largest forearm muscle bulk [66]. That is, at approximately the upper third of their forearm, as shown in Fig 2A (with the top of the armband at a median of 28% of the way down the forearm from the medial epicondyle to the ulnar styloid process). The Myo armband contained eight surface electrodes to collect EMG data at a sampling rate of 200 Hz. The Myo armband also contained one IMU to collect limb position data (three accelerometer, three gyroscope, and four quaternion data streams) at 50 Hz. Myo Connect software was used to stream and record EMG and IMU data in Matlab.

**Fig 2.**
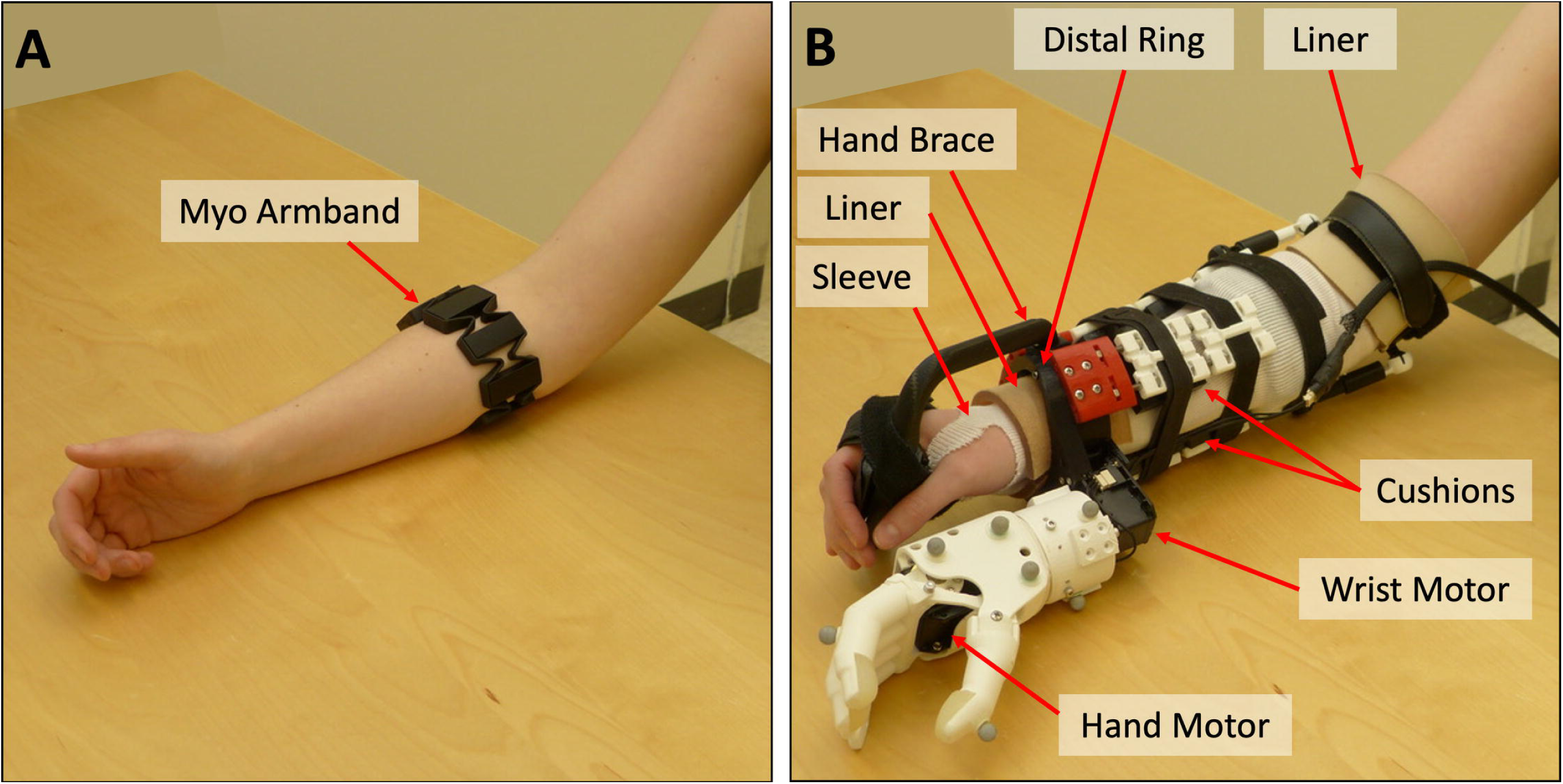
Myo armband and simulated prosthesis. A) Myo armband on a participant’s forearm and B) simulated prosthesis on a participant’s forearm, with labels indicating the sleeve, two pieces of liner, hand brace, distal ring, cushions, wrist motor, and hand motor. Adapted from Williams et al. [26]

The EMG data from the Myo armband were filtered using a high pass filter with a cutoff frequency of 20 Hz (to remove movement artifacts), as well as a notch filter at 60 Hz (to remove electrical noise). The accelerometer data streams were upsampled to 200 Hz (using previous neighbour interpolation) to align them with the corresponding EMG data. Data were then segmented into windows (160-millisecond with a 40-millisecond offset).

### Simulated Prosthesis & Donning Procedure

The simulated prosthesis used in this study was the 3D-printed Modular-Adaptable Prosthetic Platform (MAPP) [67] (shown in Fig 2B). It was fitted to each SP Participant Group member’s right arm for simulation of transradial prosthesis use. The MAPP’s previously-published design [67] was altered to improve wearer comfort in our study—the distal ring was made to resemble the oval shape of a wrist and the hand brace was elongated so that the distal ring would sit more proximally on the wearer’s wrist. A 3D-printed robotic hand [68] was affixed to the MAPP beneath the participant’s hand. Wrist rotation capabilities were also added to the device. Hand and wrist movements were each powered by a Dynamixel MX Series motor (Robotis Inc., Seoul, South Korea).

After placement of the Myo armband, each SP Participant Group member donned a thin, protective sleeve and then the simulated prosthesis. To increase participant comfort, pieces of thermoplastic elastomer liner were placed inside the distal ring and just above the participant’s elbow, and 3D-printed cushions, made of Ninjaflex Cheetah filament (Ninjatek, Inc.), were placed throughout the device socket (shown in Fig 2B). The secureness of the device and the participants’ comfort were checked before proceeding with controller training.

### Control Model Descriptions & Training Routines

#### RCNN-TL’s Model

Bayesian optimization automatically determined the number of convolution layers, number of filters, filter size, pooling size, and patience required for the classifier used in this controller. Optimization was performed in two steps: first, the number of layers along with each hyperparameter being optimized were determined using a broad range of values; thereafter, values were refined using a narrower range (centered at earlier optimized values). RCNN-TL’s model architecture consisted of 19 layers, as illustrated in Fig 3. In this model, a sequence input layer first received and normalized the training data. Then, a sequence folding layer was used, allowing convolution operations to be performed independently on each window of EMG and accelerometer data. This was followed by a block of four layers: a 2D convolution, a batch normalization, a rectified linear unit (ReLU), and an average pooling layer. This block of layers was repeated once more. Each of the two average pooling layers had a pooling size of 1×4. A block of three layers followed: a 2D convolution, a batch normalization, and a ReLU layer. The optimal number of filters in the convolution layers were determined to be 4, 16, and 32, respectively, and each had a filter window size of 1×3. The next layers included a sequence unfolding layer (to restore the sequence structure), a flatten layer, a long short-term memory (LSTM) layer, and a fully connected layer. Finally, a softmax layer and classification layer were used. To prevent overfitting, a patience parameter was set to trigger early stopping when the validation loss increased five times (similar to methods used in other works, including Côtée-Allard et al. [69]).

**Fig 3.**
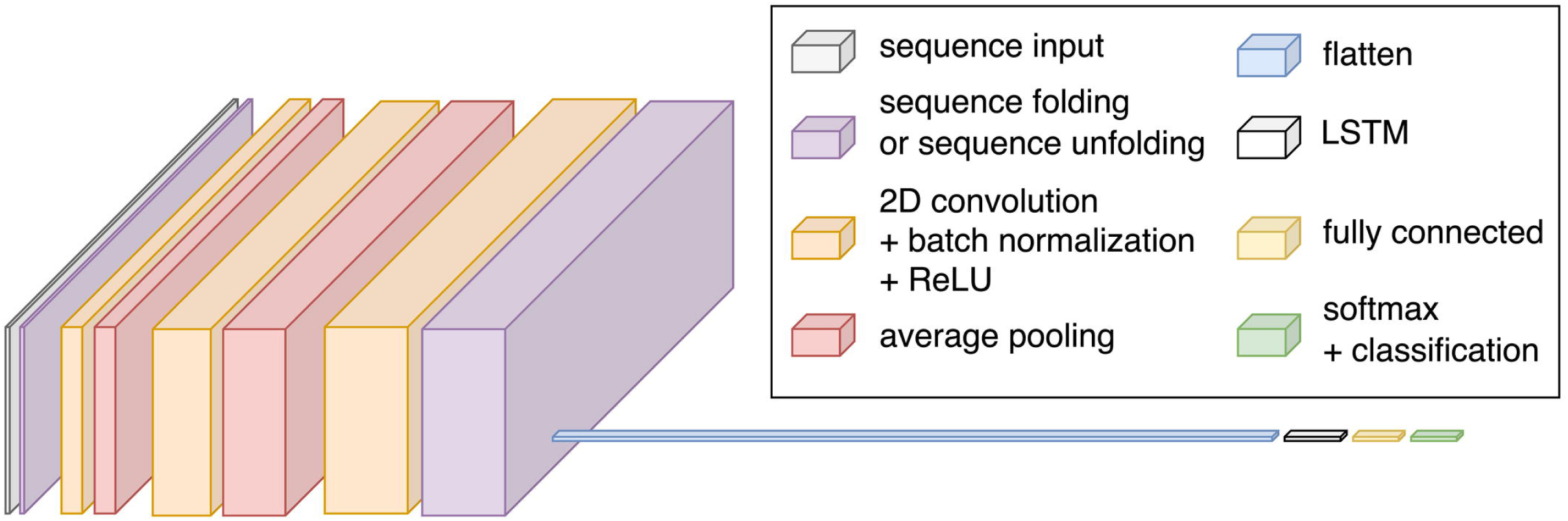
Architecture of RCNN-TL’s model: sequence input layer; sequence folding layer; two blocks of 2D convolution, batch normalization, rectified linear unit (ReLU), and average pooling; one block of 2D convolution, batch normalization, and ReLU; sequence unfolding layer; flatten layer; long short-term memory (LSTM) layer; fully connected layer; softmax layer; and classification layer. Adapted from Williams et al. [26]

#### RCNN-TL’s Model Pre-Training Routine

General Participant Group members followed onscreen instructions, performing muscle contractions in 5 wrist positions (rest, flexion, extension, pronation, and supination; shown in Fig 1A), for 5 seconds each. The muscle contractions were performed twice in 4 limb positions: arm at side, elbow bent at 90°, arm out in front at 90°, and arm up 45° from vertical (shown in Fig 1A). This position-aware routine was similar to those used in other real-time control studies aiming to mitigate the limb position effect [8, 23, 66]) and took approximately 200 seconds. The resulting EMG and accelerometer data, plus corresponding classes of muscle contractions, were used to pre-train RCNN-TL’s model.

#### RCNN-TL’s Model Retraining Routine

Our previous offline research [70] examined methods of reducing user training burden and uncovered a shortened/optimized routine that still yielded high predictive accuracy. In keeping with this, the SP Participant Group members followed onscreen instructions, performing muscle contractions in the same 5 wrist positions (shown in Fig 1B), for *only 2* (rather than 5) seconds each. The muscle contractions were performed twice in *only 3* (not 4) limb positions: arm at side, elbow bent at 90°, and arm up 45° from vertical (shown in Fig 1B). The resulting EMG and accelerometer data, plus corresponding classes of muscle contractions, were used to retrain RCNN-TL’s model.

#### LDA-Baseline’s Model

Four commonly used EMG features were chosen for implementation of this controller’s classifier: mean absolute value, waveform length, Willison amplitude, and zero crossings [71]. These features were calculated for each channel within each window of EMG data. A pseudo-linear LDA discriminant type was used, given that columns of zeros were occasionally present in some classes for some features (including Willison amplitude and zero crossings).

#### LDA-Baseline’s Model Training Routine

SP Participant Group members followed onscreen instructions, performing muscle contractions in 5 wrist positions (shown in Fig 1B), for 5 seconds each. The muscle contractions were performed twice, with the participants’ elbow bent at 90° (shown in Fig 1B). This single-position routine mimicked standard myoelectric prosthesis training [3] and took approximately 50 seconds. The resulting EMG data and corresponding classes of muscle contractions were used to train LDA-Baseline’s model.

#### RCNN-TL & LDA-Baseline Implementation

Each model was trained using Matlab software running on a computer with an Intel Core i9-10900K CPU (3.70 GHz) with 128 GB of RAM. RCNN-TL’s and LDA-Baseline’s models were retrained/trained in median times of 3.41 and 0.39 seconds, respectively. For both controllers, Matlab code was written to receive signal data and subsequently classify wrist and hand movements. Code was also written to send motor instructions, based on the resulting classifications, to brachI/Oplexus software [72] (flexion controls hand close, extension controls hand open, pronation controls wrist counter-clockwise rotation, and supination controls wrist clockwise rotation). brachI/Oplexus relayed the corresponding control signals to the simulated prosthesis’ motors. The positions of the motors were recorded with a sampling rate of 50 Hz.

### Simulated Device Control Practice & Testing Eligibility

During each testing session, SP Participant Group members took part in a control practice period (approximately 40 minutes), during which they were taught how to operate the simulated prosthesis using isometric muscle contractions, under three conditions:

1. Controlling the hand open/close while the wrist rotation function was disabled. They practiced grasping, transporting, and releasing objects at varying heights.
2. Controlling wrist rotation while the hand open/close function was disabled. They practiced rotating objects at varying heights.
3. Controlling the hand open/close function in concert with the wrist rotation function. They practiced tasks that involved grasping, transporting, rotating, and releasing objects at varying heights.

Following their practice period, participants were tested to determine whether they could reliably control the simulated prosthesis. Two cups were situated in front of them at two different heights, with a ball in one of the cups. Participants were asked to pour the ball between the two cups, and instances when they dropped the ball or a cup were recorded. If participants could not complete at least 10 pours with a success rate of at least 75% within 10 minutes, the session was ended, and they were removed from the study. Recall that one participant was removed (as stated in the Participants section), given that they could not complete this activity with LDA-Baseline in their first session.

### Motion Capture Setup & Kinematic Calibrations

For participants who were deemed eligible for controller testing, the following motion capture steps were undertaken: **Step 1: Motion Capture Setup**—An 8-camera OptiTrack Flex 13 motion capture system (Natural Point, OR, USA) was used to capture participant movements and task objects at a sampling rate of 120 Hz. Eight individual markers were placed on the simulated prosthesis hand, circled in Fig 4 (one on the thumb, one on the index finger, and the remaining six throughout the back and side of the hand to ensure reliable rigid body tracking). Rigid marker plates were also placed on each participant’s right forearm (affixed to the simulated prosthesis socket), upper arm, and thorax, in accordance with Boser et al.’s cluster-based marker model [73].

**Fig 4.**
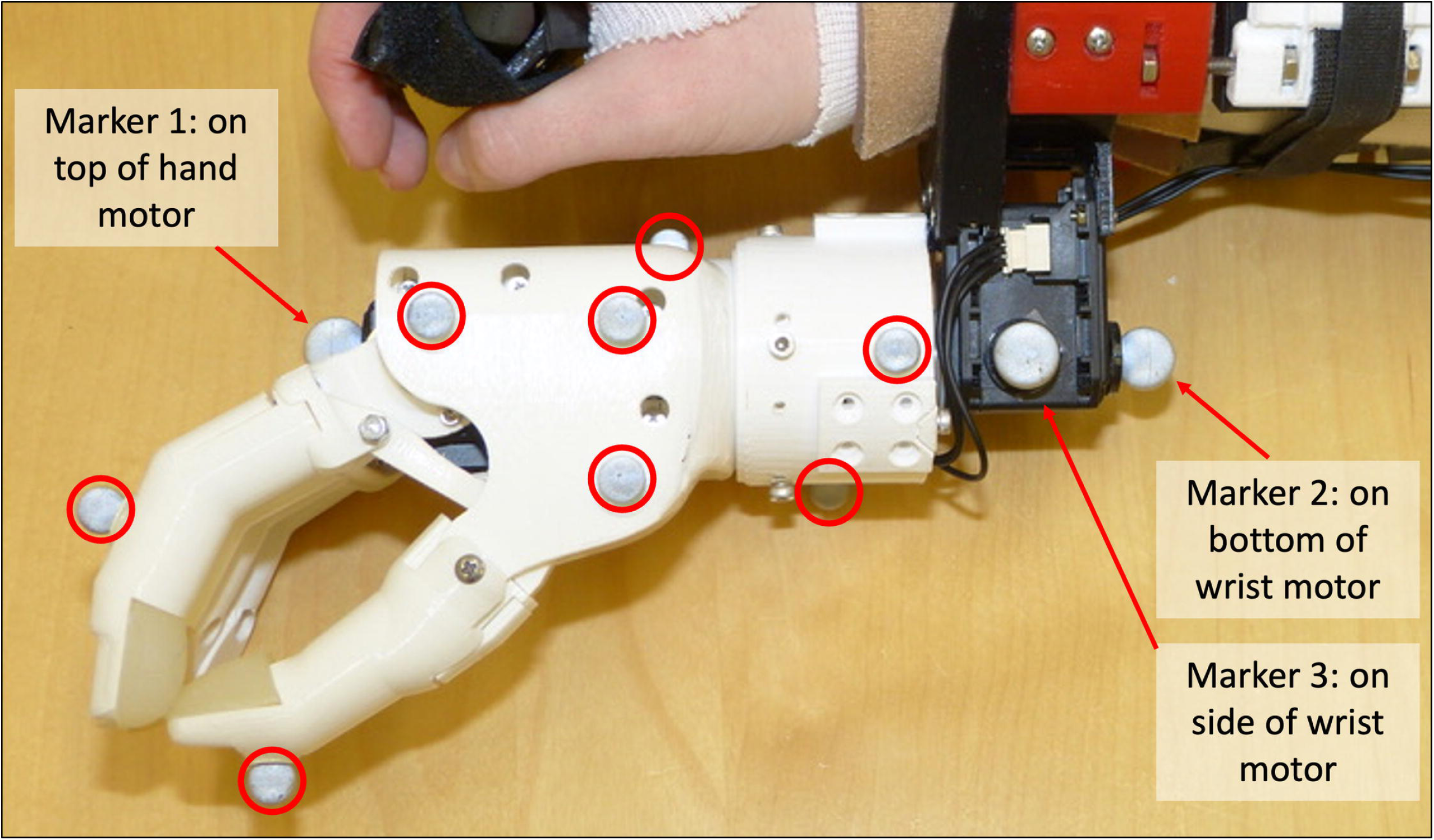
Motion capture markers affixed to the simulated prosthesis. The eight motion capture markers that remained attached to the hand are circled, and the three additional individual markers for the ski pose calibration are labelled.

#### Step 2: Kinematic Calibrations

Each participant was required to perform two kinematics calibrations. As per Boser et al., the first calibration called for participants to hold an anatomical pose [73], for capture of the relative positions of the hand markers and motion capture marker plates when wrist rotation and shoulder flexion/extension angles were at 0°. The second calibration required participants to hold a ski pose [73], for the purpose of refining wrist rotation angles. Here, three additional individual markers were affixed to the simulated prosthesis, as shown in Figure 4:

1. One marker placed on the top of the prosthesis’ hand motor, with the device hand closed
2. One marker placed on the bottom of the prosthesis’ wrist motor, forming a line with the first marker (to represent the axis about which the wrist rotation occurred)
3. One marker placed on the side of the prosthesis’ wrist motor (to create a second axis, perpendicular to the axis of wrist rotation)

Upon completion of the two kinematics calibrations, all Step 2 markers were removed. What remained were only those markers affixed during Step 1 for data collection purposes.

### Functional Tasks & Data Collection

Motion capture data were collected during the execution of the following functional tasks: **Pasta Box Task (Pasta)**—Participants were required to perform three distinct movements, where they transported a pasta box between a 1^st^, 2^nd^, and 3^rd^ location (a side table and two shelves at varying heights on a cart, including across their midline) [38]. The task setup is shown in Fig 5A. Motion capture markers were placed on all task objects, as per Valevicius et al. Participants performed a total of 10 Pasta trials. If participants dropped the pasta box, placed it incorrectly, performed an incorrect movement sequence, or hit the frame of the task cart, the trial was not analyzed. Pasta was the first of two functional tasks performed as it was considered easier.

**Fig 5.**
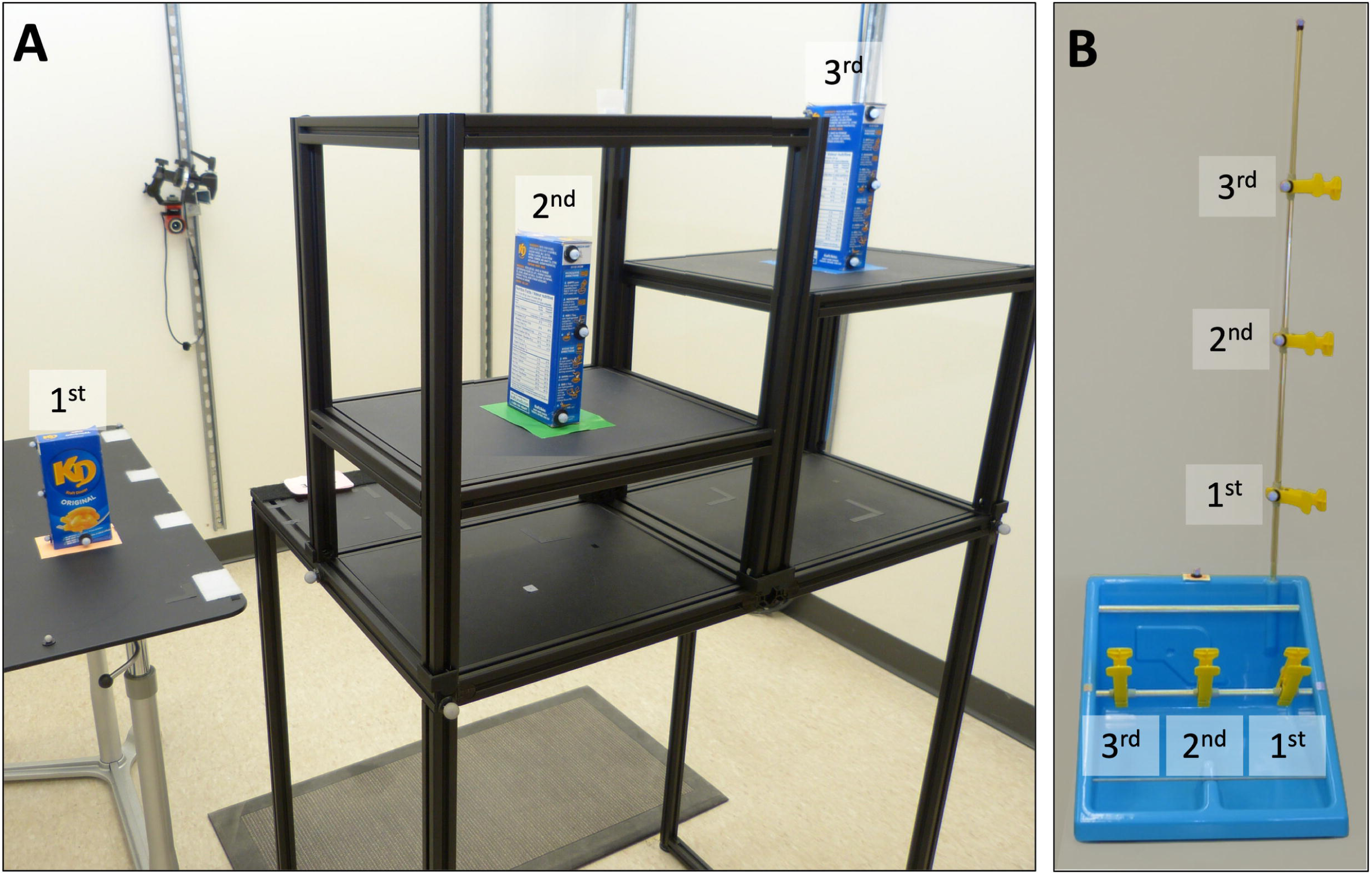
Task setup for (A) Pasta and (B) RCRT Up and Down. In panel (A), the 1^st^, 2^nd^, and 3^rd^ pasta box locations are labelled. The pasta box movement sequence is 1^st^ –*>*2^nd^ –*>*3^rd^ –*>*1^st^ locations. In panel (B), the 1^st^, 2^nd^, and 3^rd^ clothespin locations on the horizontal and vertical bars are labelled. The clothespin movement sequences in RCRT Up are horizontal 1^st^ –*>*vertical 1^st^, horizontal 2^nd^ –*>*vertical 2^nd^, and horizontal 3^rd^ –*>*vertical 3^rd^ locations. The clothespin movement sequence in RCRT Down follows the same order, but with each clothespin moved from vertical to horizontal locations.

#### RCRT

Participants were required to perform three distinct movements using clothespins. They moved three clothespins between 1^st^, 2^nd^, and 3^rd^ locations on horizontal and vertical bars [65]. To simplify trial execution, RCRT was split into RCRT Up and RCRT Down trials. The task setup for these trials is shown in Fig 5B. During Up trials, participants moved the three clothespins from the horizontal bar to the vertical bar, and during Down trials, they moved the clothespins from the vertical bar to the horizontal bar. A height adjustable cart was set such that the top of each participants’ shoulder was aligned with the midpoint between the top two targets on the vertical bar. Motion capture markers were placed on all task objects, as per our earlier research [26]. Participants performed a total of 10 Up trials and 10 Down trials. If participants dropped a clothespin, placed it incorrectly, or performed an incorrect movement sequence, the trial was not analyzed. Performance of RCRT Up and Down trials were alternated, and started with RCRT Up.

### Survey Administration

At the end of each session, each participant completed two surveys: the NASA-TLX [62] and a usability survey [64]. The former was administered using the official NASA-TLX iPad application, where participants scored their device control workload demand on a continuous rating scale with endpoint anchors of low and high. The usability survey was administered on paper, where participants marked their usability scores on a continuous rating scale with endpoint anchors of 0 and 5. In their second session, participants *were not reminded* of their survey responses from their first session.

### Data Processing & Calculation Procedures

#### Motion Capture Data Cleaning & Calculations

Motion capture marker trajectory data were cleaned and filtered. As per Valevicius et al. [38], grip aperture was calculated as the distance between the motion capture markers on the simulated prosthesis’ index and thumb, and a 3D object representing the simulated prosthesis’ hand was generated using the remaining 6 hand motion capture markers. Then, through calculations modified from Boser et al. [73], wrist rotation was calculated using the forearm and hand motion capture markers, and shoulder flexion/extension was calculated using the upper arm and thorax motion capture markers.

#### Data Segmentation

The task data were segmented in accordance with Valevicius et al. [38], as follows:

- For each *task*, the data from each *trial* were first divided into distinct *movements 1, 2, and 3* based on hand velocity and the velocity of the pasta box/clothespins during transport. **Pasta** Movements 1, 2, and 3 differentiated between: (1) reaching for the pasta box at its 1^st^ location, grasping it, transporting it to its 2^nd^ location, releasing it, and moving their hand back to a home position; (2) reaching for the pasta box at the 2^nd^ location, grasping it, transporting it to its 3^rd^ location, releasing it, and moving their hand back to the home position; and (3) reaching for the pasta box at the 3^rd^ location, grasping it, transporting it back to the 1^st^ location, releasing it, and moving their hand back to the home position. **RCRT Up** Movements 1, 2, and 3 differentiated between: (1) reaching for the 1^st^ clothespin at its 1^st^ horizontal location, grasping it, transporting it to its 1^st^vertical location, releasing it, and moving their hand back to a home position; (2) reaching for the 2^nd^ clothespin at its 2^nd^ horizontal location, grasping it, transporting it to its 2^nd^ vertical location, releasing it, and moving their hand back to the home position; and (3) reaching for the 3^rd^ clothespin at its 3^rd^ horizontal location, grasping it, transporting it to its 3^rd^ vertical location, releasing it, and moving their hand back to the home position. **RCRT Down** Movements 1, 2, and 3 differentiated between: (1) reaching for the 1^st^ clothespin at its 1^st^ vertical location, grasping it, transporting it to its 1^st^horizontal location, releasing it, and moving their hand back to a home position; (2) reaching for the 2^nd^ clothespin at its 2^nd^ vertical location, grasping it, transporting it to its 2^nd^ horizontal location, releasing it, and moving their hand back to the home position; and (3) reaching for the 3^rd^ clothespin at its 3^rd^ vertical location, grasping it, transporting it to its 3^rd^ horizontal location, releasing it, and moving their hand back to the home position.
- Then, the data from each of the three movements were further segmented into *five phases* of (1) Reach, (2) Grasp, (3) Transport, (4) Release, and (5) Home (note that the Home phase was not used for data analysis)
- Finally, *two movement segments* of (1) Reach-Grasp and (2) Transport-Release were defined for select metrics analysis
- Six final levels for data analysis resulted: *controller* (either RCNN-TL or LDA-Baseline), *task* (either Pasta, RCRT Up, or RCRT Down), *trial* (1–10), *movement* (1–3), *movement segment* (Reach-Grasp or Transport-Release), and *phase* (Reach, Grasp, Transport, or Release)

#### Grip Aperture & Wrist Rotation Re-Calculations

The grip aperture and wrist rotation angle were re-calculated using the data from the simulated prosthesis’ two motors, given that small (yet informative) adjustments in the positions of these motors may not have been detected by motion capture cameras. The positions of these motors were first upsampled to 120 Hz using linear interpolation. *Grip aperture re-calculation*: motion-capture-calculated grip aperture was used to fit a trinomial curve to transform the hand motor data to grip aperture. *Wrist motor angle re-calculation*: motion-capture-calculated wrist rotation was used to fit a binomial curve to transform the wrist motor data to wrist rotation angles.

#### Final Suite of Metrics Calculations

The final suite of metrics was calculated using the procedures described in Tables 1, 2, and 3. Note that the simultaneous wrist-shoulder movements metric was calculated only for Reach and Transport phases of RCRT Up and RCRT Down trials, because these were the only phases that required the participant to rotate the device wrist while moving their arm to a different height.

### Statistical Analysis

To investigate task performance difference between RCNN-TL and LDA-Baseline, the following statistical analyses were performed:

#### For metrics that were analyzed at the phase or movement segment level

Participants’ results were first averaged across trials and movements. If results were normally-distributed, a two-factor repeated-measures analysis of variance (RMANOVA) was conducted using the factors of controller and phase/movement segment. When the resulting controller effects or controller-phase/movement segment interactions were deemed significant (that is, when the Greenhouse-Geisser corrected p value was less than 0.05), pairwise comparisons between the controllers were conducted. If results were not normally-distributed, the Friedman test was conducted. When the resulting p value was less than 0.05, pairwise comparisons between the controllers were conducted. Pairwise comparisons (t-test/Wilcoxon sign rank test) were deemed significant if the p value was less than 0.05.

#### For metrics that were analyzed at the trial level

Participants’ results were first averaged across trials, then pairwise comparisons were conducted as detailed above.

#### For metrics that were analyzed at the task or controller level

Pairwise comparisons were conducted as detailed above.

### Limb Position Effect Identification

The limb position effect has been shown to cause control accuracy degradation and large between-participant control variation in offline research [41]. However, earlier works have not pinpointed specific instances of the effect in functional task execution data. Using the novel control characteristics metrics introduced in this work, identification of such instances is possible—larger medians and/or larger interquartile ranges (IQRs) can provide evidence of degraded control. To identify the limb position effect in this study, metrics’ medians and IQRs for Reach, Grasp, Transport, and Release phases were considered separately across movements 1, 2, and 3 of Pasta, RCRT Up and RCRT Down. An occurrence where movement variation was *not* due to the limb position effect is illustrated in Fig 6A, where the number of wrist rotation adjustments metric in RCRT Down Release phases have medians and IQRs that remain relatively constant at different limb positions. Conversely, an occurrence where movement variation *was* due to the limb position effect can be seen in Fig 6B. Here, the same metric in RCRT Down Grasp phases shows its medians and IQRs both increasing as the limb position changed.

**Fig 6.**
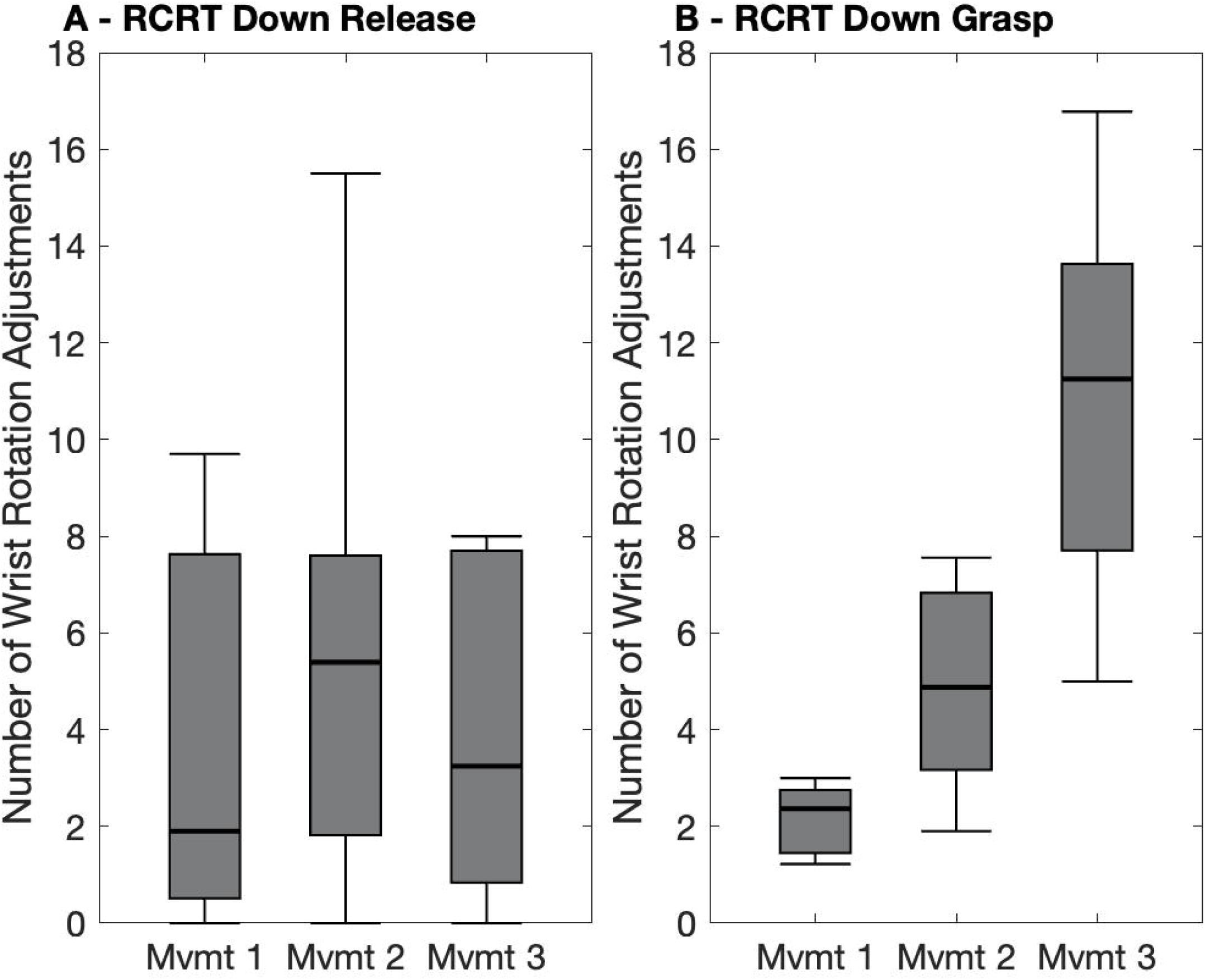
Box plots indicating LDA-Baseline number of wrist rotation adjustments in each (A) RCRT Down Release and (B) RCRT Down Grasp of each task movement (Mvmt). Medians are indicated with thick lines, and interquartile ranges are indicated with boxes.

The following limb position identification process was used to examine all control characteristics metrics for Reach, Grasp, Transport, and Release phases across the three movements of Pasta, RCRT Up, and RCRT Down:

- First, the three medians were rescaled as percentages of the maximum of the three medians
- Next, the three IQRs were rescaled as percentages of the maximum of the three IQRs
- Then, the limb position effect identification rules outlined in Table 4 were developed—through iterative trial-and-error comparisons of potential rules to visual representations of metrics’ medians and IQRs (as in Figs 6BA, B)
- The resulting rules were subsequently used to identify instances of the limb position effect. Note that for Pasta, two rule options were used, given that the limb position effect was most likely to be present in that task’s movements 2 or 3 (at the highest shelf location). For RCRT Up and RCRT Down, only one rule option was necessary, given that the limb position effect was most likely to occur in movement 3 (at the top clothespin location). For each rule, the limb position effect was identified only in instances where all conditions were met (that is, when all rules in a row of Table 4 were true).

**Table 4.** Limb position effect identification rules for control characteristics metrics. The rules used for limb position effect identification are based on each movement’s median and interquartile range (IQR). In each row, all conditions had to be true for a positive identification of the limb position effect.

| Limb Position Effect Identification Rule | Conditions Identifying Limb Position Effect |  |  |  |  |  |
| --- | --- | --- | --- | --- | --- | --- |
|  | Movement 1 Median | Movement 2 Median | Movement 3 Median | Movement 1 IQR | Movement 2 IQR | Movement 3 IQR |
| Pasta Reach and Grasp Rule | < 90% | < 75% | = 100% |  | < 55% | = 100% |
| Pasta Transport and Release Rule |  | = 100% | < 85% |  | = 100% |  |
| RCRT Up and RCRT Down Rule | < 55% | < 85% | = 100% | < 25% |  | > 80 % |

The abovementioned rules are applicable to metrics where smaller values are indicative of control improvements, as was the case with most control characteristics metrics in this work. The exception was the simultaneous wrist-shoulder movements metric, where *larger* values were indicative of improved control. To adjust for this exception, the three movements’ medians were modified by subtracting each from 100% (therefore changing these medians to represent the percent of the phase in which simultaneous movements did *not* occur). After this adjustment, the limb position identification process could be followed.

## Results

### Task Performance

The significant differences across the task performance metrics are reported in Table 5. Task specific outcomes derived from the table include:

**Table 5.**
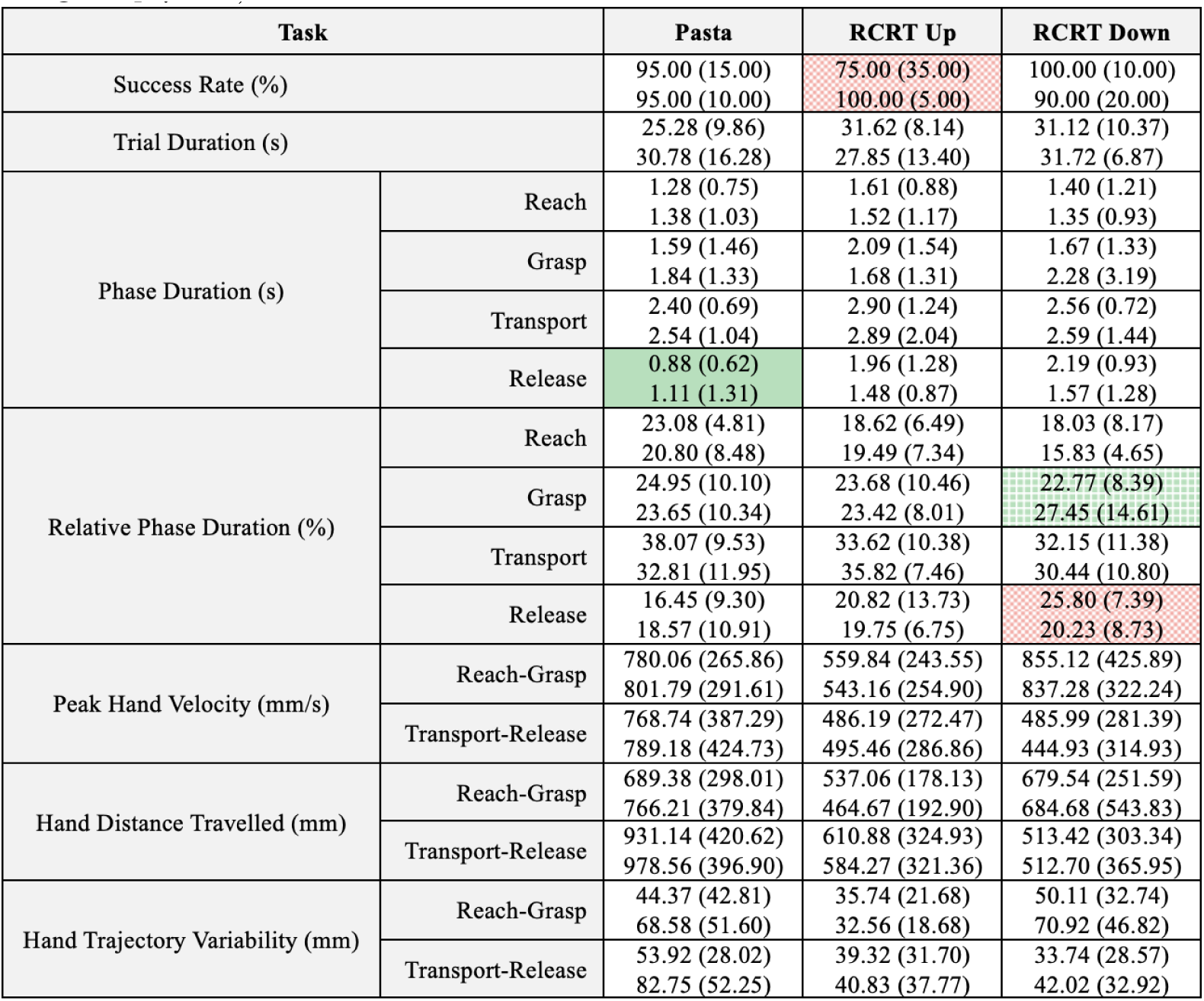
Task Performance metrics results. Each cell contains the RCNN-TL median (and interquartile range in parentheses) in the first line and the LDA-Baseline median (interquartile range) in the second line. Green cells indicate metrics in which RCNN-TL performed significantly better than LDA-Baseline (solid green: p ¡ 0.005, dense green grid: p ¡ 0.01). Red cells indicate metrics in which LDA-Baseline performed significantly better than RCNN-TL (sparse red grid: p ¡ 0.05).

#### Pasta

RCNN-TL performed significantly better than LDA-Baseline in *one* metric: Release phase duration.

#### RCRT Up

LDA-Baseline performed significantly better than RCNN-TL in *one* metric: success rate.

#### RCRT Down

RCNN-TL performed significantly better than LDA-Baseline in *one* metric: Grasp relative phase duration. LDA-Baseline performed significantly better than RCNN-TL in *one* metric: release relative phase duration.

#### Summary

Only 4 of the 48 Task Performance metrics showed significant differences, 2 of which demonstrated that RCNN-TL performed better than LDA-Baseline. It appears that a richer set of metrics is needed to better understand such outcomes—beyond those derived from task performance metrics alone.

### Control Characteristics

The significant differences across control characteristics metrics are reported in Table 6. Task specific outcomes derived from the table include:

**Table 6.**
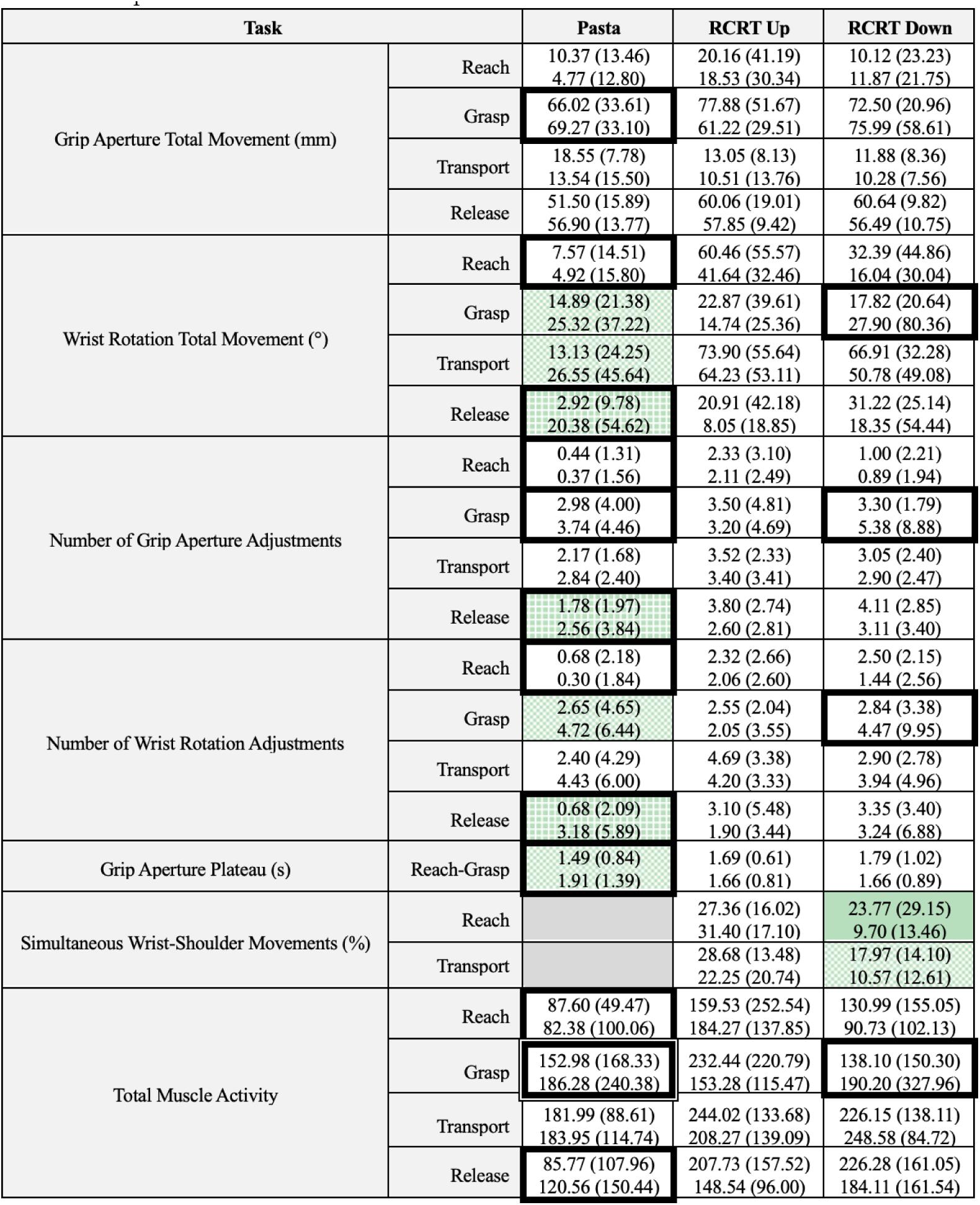
Control Characteristics metrics results. Each cell contains the RCNN-TL median (and interquartile range in parentheses) in the first line and the LDA-Baseline median (interquartile range) in the second line. Green cells indicate metrics in which RCNN-TL performed significantly better than LDA-Baseline (solid green: p ¡ 0.005, dense green grid: p ¡ 0.01, sparse green grid: p ¡ 0.05). Dark grey cells indicate instances in which a metric was not relevant. Dark cell borders indicate metrics that displayed evidence of the limb position effect under LDA-Baseline control. A double cell border indicates the metric that displayed evidence of the limb position effect under both LDA-Baseline and RCNN-TL control.

#### Pasta

RCNN-TL performed significantly better than LDA-Baseline in *seven* metrics. Examples of one such metric are illustrated in S1 Table. (for Reach-Grasp grip aperture plateau). The limb position effect was identified in *12* metrics under LDA-Baseline control, and in one metric (Grasp total muscle activity) under RCNN-TL control. Four of the seven significant RCNN-TL versus LDA-Baseline differences were in metrics that showed evidence of the limb position effect.

#### RCRT Up

No significant differences were identified, and no metrics showed evidence of the limb position effect.

#### RCRT Down

RCNN-TL performed significantly better than LDA-Baseline in *two* metrics. Examples of one such metric are illustrated in S1 Table. (for Reach simultaneous wrist-shoulder movements). The limb position effect was identified in *four* other metrics, with one such instance illustrated in Fig 6B (Grasp number of wrist rotation adjustments).

#### Summary

9 of the 81 Control Characteristics metrics showed significant differences, all of which demonstrated that RCNN-TL performed better than LDA-Baseline. Furthermore, 16 metrics showed evidence of the limb position effect. All such metrics were only identified in Pasta and RCRT Down, suggesting that these outcomes are likely influenced by the position-aware nature of RCNN-TL control.

### User Experience

User experience metrics were calculated at the controller level, rather than for each task (detailed in Table 3). There were no significant differences between RCNN-TL and LDA-Baseline. Box plots illustrating the median controller-level scores across participants can be found in Fig 7.

**Fig 7.**
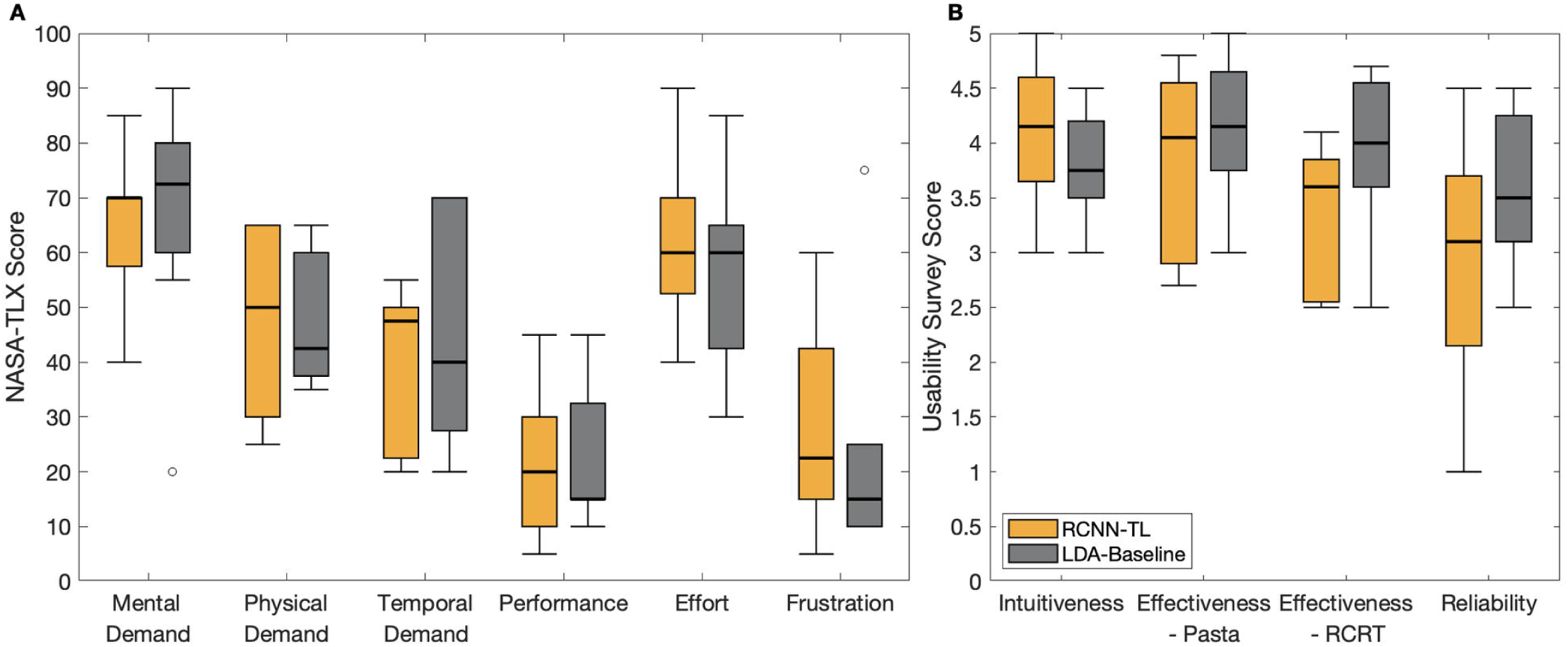
Box plots indicating user experience metrics results with RCNN-TL (orange) and LDA-Baseline (grey) for: (A) NASA-TLX, and (B) usability survey. Medians are indicated with thick lines, interquartile ranges are indicated with boxes, and outliers are indicated with circles.

Of note, RCNN-TL scored better than LDA-Baseline in the NASA-TLX’s Mental Demand dimension and in the usability survey’s Intuitiveness dimension. These results suggest that RCNN-TL offered more intuitive control. The two controllers had equal median scores in NASA-TLX’s Effort dimension. LDA-Baseline scored better than RCNN-TL in all other dimensions.

## Discussion

The suite of myoelectric prosthesis control evaluation metrics introduced in this work (detailed in Tables 1, 2, and 3) yielded informative limb position effect-related outcomes that could only be speculated upon in our earlier work [26]. What follows is a discussion about the metrics-driven findings from this current work, with a focus on understanding *when* and *why* limb position variations caused control challenges during participants’ execution of the Pasta Box Task (Pasta) and the Refined Clothespin Relocation Test (RCRT Up and RCRT Down).

### Findings from Control Characteristics Metrics

#### Limb Position Effect Identification

To our knowledge, no other study has identified occurrences of the limb position effect using functional task assessment outcomes. In this work, we used functional tasks to assess device control and found that analysis of our control characteristics metrics *did* uncover instances of the limb position effect. In LDA-Baseline results, instances were identified in Pasta and RCRT Down (16 of 81 cells with dark borders in Table 6), and never in RCRT Up. Note that for RCNN-TL, one instance of the limb position effect was uncovered—Grasp total muscle activity in Pasta. However, this instance may simply be due to inevitable positional EMG signal variations, rather than due to control degradation. Based *solely* on LDA-Baseline results, we surmised the following:

- Raised arm positions in the sagittal plane caused grasp challenges for participants, as evidenced by the identification of the limb position effect only during the Grasp phases of RCRT Down (in four metrics).
- Raised arm positions in the sagittal plane did *not* cause release challenges for participants, as evidenced by the *absence* of limb position effect identification in RCRT Up. Logically, as hand opening during object release phases is controlled by wrist extension muscle activation, classification of wrist extension was not affected by the limb position effect.
- Arm movements along the frontal plane caused further control challenges for participants. Not only did Pasta require participants to perform arm raises in the sagittal plane, large cross-body and away-from-body movements had to be introduced to accomplish this task. The limb position effect was detected in three of four Pasta phases (four times in Reach, once in Reach-Grasp, three times in Grasp, and four times in Release)

So overall, LDA-Baseline control often appeared to be impeded by shoulder position fluctuations in the frontal plane. Furthermore, arm raises limited to the sagittal plane caused only grasp control deterioration for this controller. Both such circumstances identify catalysts for limb position effect control challenges.

#### Evidence of Limb Position Effect Mitigation

Recall that Table 6 also identified significant differences in control characteristics metrics, where green cells indicated instances where RCNN-TL performed significantly better than LDA-Baseline. Nine of the 81 metrics (cells) were significant (shaded in green), indicating that RCNN-TL always performed the same as, or significantly better than, LDA-Baseline for these metrics. Furthermore, all such significant differences presented in Table 6 occurred in Pasta and RCRT Down, and never in RCRT Up. This coincides with those tasks where instances of the limb position effect were identified, suggesting that RCNN-TL successfully mitigated such occurrences.

Interestingly, RCNN-TL performed significantly better than LDA-Baseline in several Pasta metrics, even though Pasta involves numerous limb positions that were not included in RCNN-TL’s pre-training/retraining routines. We speculate that the pre-training data from 19 individuals provided sufficient variety to result in a controller that is robust to limb positions not included in its training routines.

Significant control characteristics differences between RCNN-TL and LDA-Baseline were not identified for RCRT Up. Recall that the problem was *not* identified in RCRT Up, despite this task’s requirement for varied limb positions. So, if the limb position effect did not cause control degradation for either controller, then perhaps: (a) LDA-Baseline simply performed well during this task and control improvements were not necessary, or (b) RCNN-TL control should be improved in instances when the limb position effect is *not* evident.

#### Merits of Control Characteristics Metrics

Significant differences between RCNN-TL and LDA-Baseline were identified in at least one phase/movement segment for all control characteristics metrics analyzed, with the exceptions of total grip aperture movement and total muscle activity. Still, these two metric exceptions might yield outcomes beneficial to other controller comparisons and should not be discounted from the metrics introduced in this work. Total grip aperture movement, for instance, might help to identify grasping efficiency during task execution, and total muscle activity might help to identify muscle exertion required for task completion. Future controller comparisons are expected to determine whether these metrics are sensitive to controller variations.

### Findings from Task Performance Metrics

Table 5 identified 2 of 48 metrics that showed RCNN-TL performing significantly better than LDA-Baseline (for Release phase duration in Pasta and Grasp relative phase duration in RCRT Down), and 2 of 48 metrics that showed the contrary (for success rate in RCRT Up and Release relative phase duration in RCRT Down). These outcomes coincided with those of our earlier work [26], however, the control characteristics metrics introduced in this study facilitated a deeper understanding of why task performance deteriorated at times—specifically, when instances of the limb position effect hampered control. The following task performance insights were uncovered in this work:

- RCNN-TL successfully mitigated the limb position effect, as evidenced by the two specific instances when its control was significantly better than that of LDA-Baseline—in Pasta Release and RCRT Down Grasp phases. Our control characteristics analysis revealed that participants struggled during these phases (identified as instances where the limb position effect occurred). Such struggles were apparent when participants used LDA-Baseline, but not so when using RCNN-TL. So, RCNN-TL likely remedied control degradation introduced by the limb position effect.
- RCNN-TL may not have performed well in instances where the limb position effect was *not* evidenced. Consider that two task performance metrics showed that RCNN-TL performed significantly *worse* than LDA-Baseline: (1) the relative duration of the RCRT Down Release phases, and (2) the RCRT Up success rate. Furthermore, consider that analysis of control characteristics revealed that device control was *not* affected by the limb position effect in either the RCRT Down Release phase, or any phase of RCRT Up. Together, these outcomes support the hypothesis that RCNN-TL may not have performed well in instances where the limb position effect was *not* evidenced.

**–** Of note, the lower RCRT Up success rate was due to clothespins being dropped by participants. This tendency towards unintended hand opening is in keeping with RCNN-TL’s control characteristics results versus those of LDA-Baseline, with higher medians in grip aperture total movement, higher medians in number of grip aperture adjustments, along with larger IQRs in Reach and Grasp phases of these same metrics.

### Findings from User Experience Metrics

Fig 7 presented the user experience metrics. Despite this work’s improved control outcomes, no significant differences were identified in the NASA-TLX and usability surveys. This work uncovered the following insights, to guide future use of user experience metrics:

- Without the provision of participants’ scores from their first testing session upon return for their second session (following the washout period of at least one week), their initial anchor scores of “good” and “poor” were not likely to have been precisely recalled.
- Given that participants were without limb loss, they only had a perception of fully functional control using their intact hand and wrist. They did not have a baseline perception of poor or diminished control, as none had prior experience with a simulated prosthesis. As a result, their subjective anchor scores of “good” and “poor” in their first session were likely influenced by their perception of perfect control, whereas in their second session, they might have been further influenced by their first session’s simulated prosthesis control experience.

Overall, a specific question about which controller each participant preferred (asked at the end of their second session) would better gauge their controller partiality. In addition, reminding participants of their first-session scores immediately prior to their second session survey completion, might address expectation-related variability in anchor scores [74]. After this current study was conducted, a new Prosthesis Task Load Index (PROS-TLX) was developed and validated [75], and should be considered in future comparative prosthesis control research.

### Evidence of Training Routine Reduction

The General Participant Group performed a long (200-second) pre-training routine prior to RCNN-TL use. This pre-training duration is similar to that of position-aware controller solutions in the literature [8, 12, 20, 66, 76–78]. RCNN-TL retraining, as performed by the SP Participant Group, was accomplished using a shortened (60-second) routine. That is, a 70% decrease in model training duration resulted due to the introduction of transfer learning. This current research, therefore, confirms that TL *is* a valuable adjunct to RCNN-based classification control, as it offers a model starting point that needs only to be calibrated using a smaller amount of individual-specific data. Notably, a TL solution is not possible with LDA-based control. Overall, this training routine reduction solution shows promise towards solving the limb position effect challenge—*without* the requirement of a burdensome training routine.

### Limitations

As a first limitation of this study, participants without upper limb impairment were recruited rather than myoelectric prosthesis users. Although these participants learned how to control a simulated prosthesis, further practice may have been necessary to accurately represent the control capabilities of myoelectric prosthesis users. Secondly, the implementation of the surveys may not have adequately captured user experience data. Thirdly, previous work had suggested that the conditions under which the RCNN-TL pre-training data was collected could result in control flaws [26]—because pre-training data from participants *without* a donned simulated prosthesis were too dissimilar from those *with* a donned simulated prosthesis (the conditions under which the retraining and device use occurred). This testing condition dissimilarity consideration was not examined in the current study. Finally, although our experimentation used optical motion capture technology to gather movement data, our suite of control metrics is not reliant on this data for metrics calculations. Alternative sources that capture grip aperture and wrist rotation (such as the positions of device motors), along with shoulder flexion/extension (such as IMUs) can be used for such calculations. Furthermore, other methods of segmenting functional tasks into Reach, Grasp, Transport, and Release phases (such as IMUs) c

### Future Work

#### RCNN-TL future work

Next steps for RCNN-TL should focus on improvements to device control in instances when the limb position effect is least likely to occur.

Improvements to pre-training data collection conditions need to be studied. Examination of RCNN-TL using myoelectric prosthesis users is also a necessary step.

#### Suite of control evaluation metrics future work

To verify whether the suite of metrics introduced in this study are beneficial, future work should examine different controllers, using both participants without limb loss wearing a simulated prosthesis and actual myoelectric prosthesis users.

## Conclusion

This work reinvestigated earlier comparative RCNN-TL versus LDA-Baseline research, which recommended that pattern recognition-based control not be judged by *task performance* alone, but rather, that *control characteristics* also be measured [26]. Then collectively, the task performance and control characteristics should be weighed against qualitative *user experience* [26]. The current study heeded these recommendations, and in doing so, contributed and tested a viable suite of myoelectric prosthesis control evaluation metrics for use in future comparative control model research. Using these metrics, this study has contributed insights into occurrences and implications of the limb position effect challenge and offered validation that TL-based neural network control solutions show promise towards solving this pervasive problem. The suite of metrics introduced and subsequently used in this work is expected to benefit future research intent on improving rehabilitation device control.

## Supporting information

**S1 Table. Metrics examples.** This table contains figures that exemplify “good” and “poor” results for selected metrics. For each metric presented, a description of the example figure precedes that associated good/poor graphs.

## Acknowledgments

We thank Quinn Boser, Thomas R. Dawson, Michael R. Dawson, and Albert Vette for experimental design and data processing assistance.

## Notes

### Competing Interest Statement

The authors have declared no competing interest.

