## Supplemental Table 1 for "A multifaceted suite of metrics for comparative myoelectric prosthesis controller research"

### S1 Table. Metrics examples.

This table contains figures that exemplify “good” and “poor” results for selected metrics. For each metric presented, a description of the example figure precedes that associated good/poor graphs.

<table>
<tr><th></th><th>Poor Result Example</th><th>Good Result Example</th></tr>
<tr><td rowspan="3">Hand Trajectory Variability</td><td colspan="2">The black lines illustrate a given participant's hand movement paths throughout the movement 3 Reach-Grasp movement segments of Pasta in each trial. The blue shaded area illustrates the standard deviations at each sampled point. Both figures were generated using data from the same participant, with the poor result example illustrating a trial in which they were using the LDA-Baseline and the good result example illustrating a trial in which they were using the RCNN-TL-Class. The hand trajectory variability calculated in these examples are indicated in each figure.</td></tr>
<tr><td>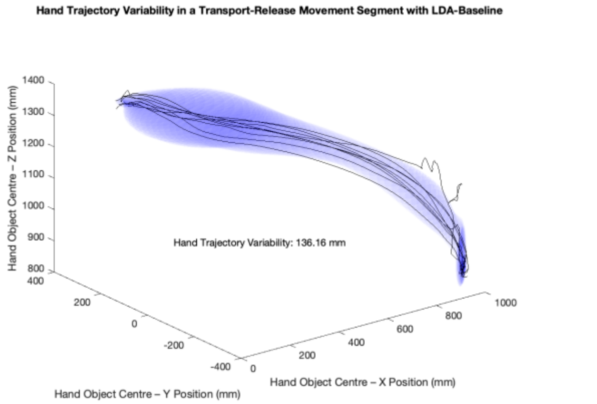
</td><td>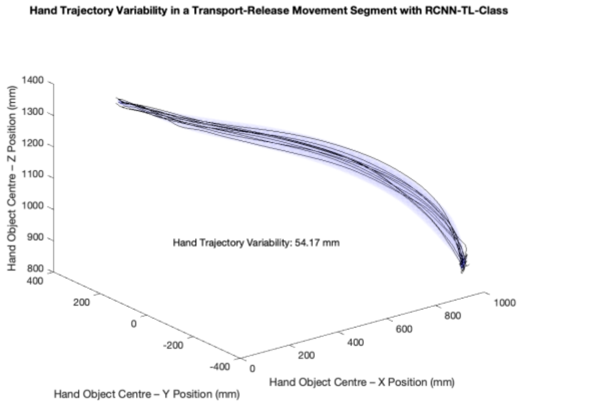
</td></tr>
<tr><td>The poor result example illustrates a large standard deviation, resulting in a large hand trajectory variability (136.16 mm).</td><td>The good result example illustrates a smaller standard deviation, resulting in a smaller hand trajectory variability (54.17 mm).</td></tr>
<tr><td rowspan="3">Total Grip Aperture Movement</td><td colspan="2">The blue line illustrates the grip aperture throughout a movement 3 Grasp phase of Pasta. Both figures were generated using data from the same participant, with the poor result example illustrating a trial in which they were using the LDA-Baseline and the good result example illustrating a trial in which they were using the RCNN-TL-Class. The total grip aperture movement are presented in each figure.</td></tr>
<tr><td>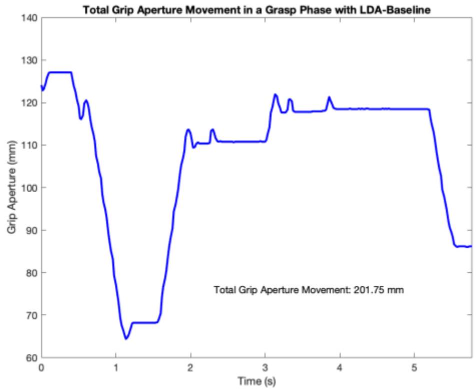
</td><td>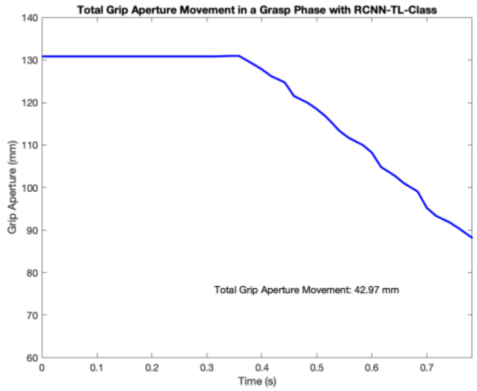
</td></tr>
<tr><td>The poor result example illustrates hand openings throughout the grasp phase, resulting in a large total grip aperture movement (201.75 mm).</td><td>The good result example illustrates a grip aperture movement with less fluctuation. This resulted in a smaller total grip aperture movement (42.97 mm).</td></tr>
</table>

<table>
<tr><th></th><th>Poor Result Example</th><th>Good Result Example</th></tr>
<tr><td rowspan="2">Number of Grip Aperture Adjustments</td><td colspan="2">The blue line illustrates the filtered grip aperture throughout a movement 3 Grasp phase of Pasta. The dashed vertical lines indicate grip aperture adjustments that were detected. Both figures were generated using data from the same participant, with the poor result example illustrating a trial in which they were using the LDA-Baseline and the good result example illustrating a trial in which they were using the RCNN-TL-Class. The number of grip aperture adjustments are presented in each figure.</td></tr>
<tr><td>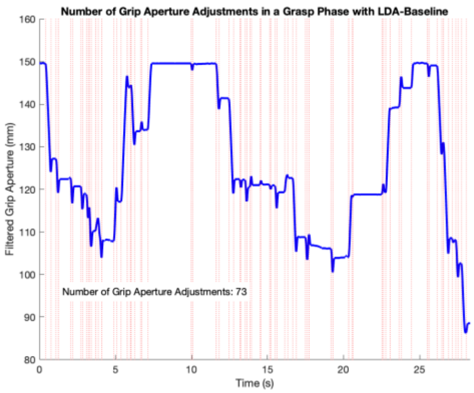

The poor result example identifies several adjustments (73).</td><td>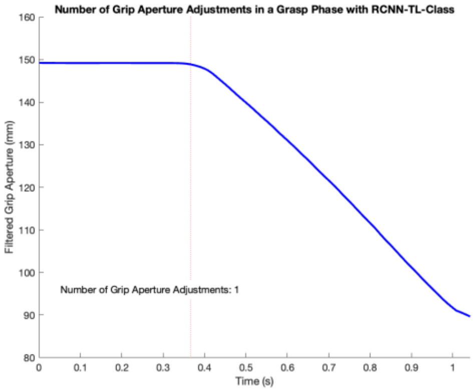

The good result example identifies only 1 adjustment, at which the hand first started to close.</td></tr>
<tr><td rowspan="2">Grip Aperture Plateau</td><td colspan="2">The blue line illustrates the grip aperture throughout a movement 3 Reach-Grasp movement segment of Pasta. The shaded area indicates the portions of the movement segment in which the grip aperture plateau requirements were met. Both figures were generated using data from the same participant, with the poor result example illustrating a trial in which they were using the LDA-Baseline and the good result example illustrating a trial in which they were using the RCNN-TL-Class. Note that the same x axis scale was used in both figures to further illustrate the grip aperture plateau. The grip aperture plateau time is presented in each figure.</td></tr>
<tr><td>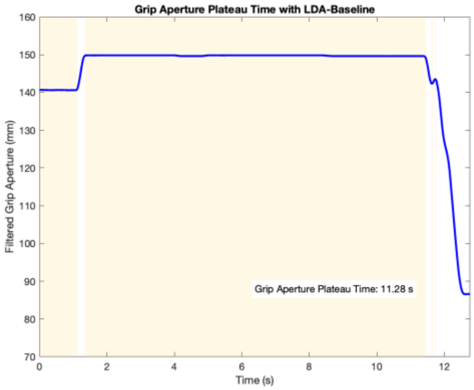

The poor result example identifies a long grip aperture plateau time (11.28 s).</td><td>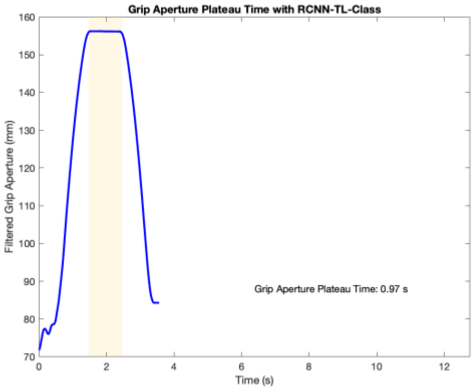

The good result example identifies a short grip aperture plateau time (0.97 s).</td></tr>
</table>

<table>
<tr><th></th><th>Poor Result Example</th><th>Good Result Example</th></tr>
<tr><td rowspan="2">Wrist-Shoulder Simultaneous Movements</td><td colspan="2">The solid blue line illustrates the filtered absolute wrist rotation angular velocity throughout a movement 3 Reach phase of RCRT down, and the solid magenta line illustrates the filtered absolute shoulder flexion/extension angular velocity throughout that same phase. The dashed horizontal blue and magenta lines indicate the thresholds corresponding to these angular velocities. Finally, the shaded areas indicate the portions of the phase in which both angular velocities were above their respective thresholds. The figures were generated using data from the same participant, with the poor result example illustrating a trial in which they were using the LDA-Baseline and the good result example illustrating a trial in which they were using the RCNN-TL-Class. The wrist-shoulder simultaneous movements are indicated in each figure.</td></tr>
<tr><td>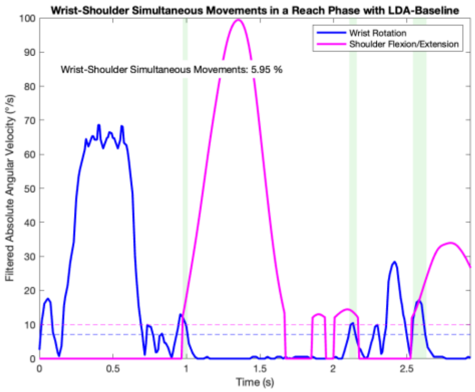<br><br>The poor result example identifies wrist rotation control when the shoulder is usually not moving, resulting in a small wrist-shoulder simultaneous movement (5.95 %).</td><td>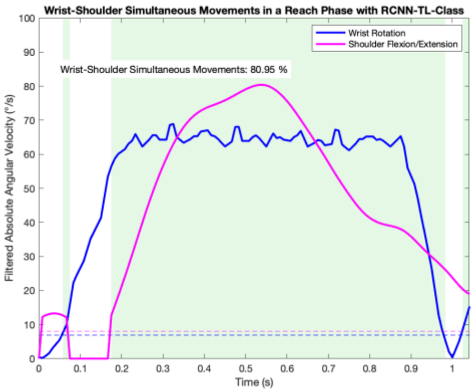<br><br>The good result example identifies wrist rotation control during shoulder movements, resulting in a large wrist-shoulder simultaneous movement (80.95%).</td></tr>
<tr><td rowspan="2">Total Muscle Activity</td><td colspan="2">The 8 coloured lines illustrate the 8 EMG signals throughout a movement 3 Grasp phase of Pasta. The black line illustrates the Grip Aperture throughout that same phase, to aid in the understanding of the EMG signals. Both figures were generated using data from the same participant, with the poor result example illustrating a trial in which they were using the LDA-Baseline and the good result example illustrating a trial in which they were using the RCNN-TL-Class. Note that the same x axis scale was used in both figures to further illustrate the differences in time required to close the hand. The total muscle activity is indicated in each figure.</td></tr>
<tr><td>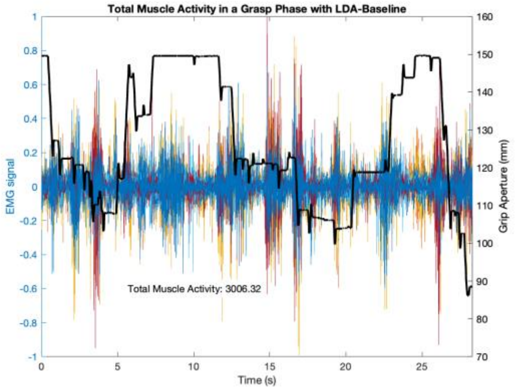<br><br>The poor result example identifies difficulties in closing the hand, resulting in a long phase and strong EMG signals. So, a large total muscle activity resulted (3006.32).</td><td>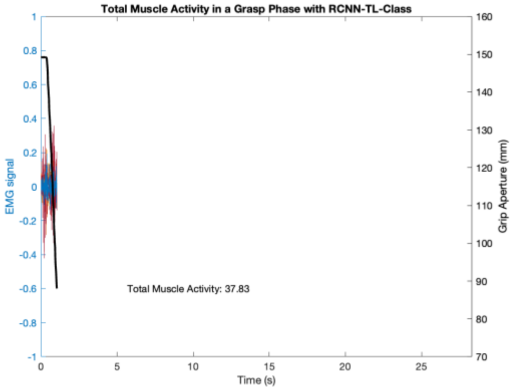<br><br>The good result example identifies more ease in closing the hand, resulting in a shorter phase and weaker EMG signals. So, a smaller total muscle activity resulted (37.83).</td></tr>
</table>
